# MedImg: a database for public medical image integration

**DOI:** 10.1101/2024.04.16.589768

**Authors:** Bitao Zhong, Rui Fan, Xiangwen Ji, Qinghua Cui, Chunmei Cui

## Abstract

The advancements of deep learning algorithms in medical image analysis has garnered tremendous attention in recent years. Several studies have reported that the models have achieved and even surpassed human performance, whereas the translation of these models into clinical practice is still accompanied by various challenges. A major challenge is the large-scale and well characterized dataset to validate the generalization of models. Therefore, we collected diverse medical image datasets from multiple public sources containing 103 datasets, 1,622,956 images. These images are derived from 14 modalities like XR, CT, MRI, OCT, ultrasound, and endoscopy, and from 9 organs such as lung, brain, eye, and heart. Subsequently, we constructed an online database, MedImg, which incorporates and hierarchically organizes medical images to facilitate data access. MedImg serves as an intuitive and open-access platform for contributing to deep learning-based medical image analysis, accessible at https://www.cuilab.cn/medimg/.

## Introduction

Medical imaging is an indispensable means to localize lesions and aid in the diagnosis and treatment of diseases [1], in which image interpretation to draw clinical conclusions is typically carried out by physicians. The computer-assisted diagnosis system emerges to expedite the diagnosis process and to reduce the false positive/negative resulting from variations in expertise [2]. With the vast advantage for automated feature learning and extraordinary performance, deep learning has become widely popular in the field of medical image analysis, including image classification for differentiating disease and normal individuals [3-5], organ or lesion detection aiming at identifying the small lesion region on a full image [6, 7], image segmentation for partitioning an image into multiple segments to perform localization or quantification analysis [8-10], and registration for aligning more images across modalities or time into one coordinate system [11, 12]. One of clinical applications achieving good performance is OSAIRIS, precisely segmenting the cancer region from the healthy organ, which revolutionizes the preparation of scans and greatly reduces waiting time of patients between referral and treatment initiation. Despite the explosion of studies focused on applying deep learning algorithms to medical images, transferring these models into clinical practices remains a challenge [13]. One of primary obstacles is the lack of large quantities of available image data, which is of great importance to optimally train, validate, and test the algorithms.

Currently, several databases gathering a wealth of medical images have been presented. For example, the Cancer Imaging Archive Collections (TCIA) [14] shares more than 30 million radiology images of cancers from around 37,568 subjects, organized by National Cancer Institute (NCI); the Open Access Series of Imaging Studies (OASIS) [15, 16] platform includes abundant neuroimaging datasets with multiple modalities, covering 3,059 subjects; The Alzheimer’s Disease Neuroimaging Initiative (ADNI, http://www.adni.loni.usc.edu) database collects magnetic resonance imaging (MRI) and positron emission tomography (PET) images from over 1700 individuals with cognitive impairment or AD; the National COVID-19 Chest Imaging Database (NCCID) [17] consists diverse chest images from over 7,000 patients, together with detailed clinical information, which is built for improving the healthcare delivery for COVID-19 patients. Furthermore, there are a fraction of artificial intelligence (AI) algorithms-related competitions publishing large image datasets, for example, musculoskeletal radiographs (MURA) containing 40,561 multi-view radiographic X-Ray (XR) images from 14,863 studies and 12,173 patients [18]. However, we observed that these datasets or databases focused mainly on unitary organ/disease or single imaging modality hindering the advancement of general deep learning models. It is quite necessary to develop a comprehensive platform with abundant medical images across modalities, organs, and geographic areas.

For this purpose, we proposed an online medical image database, named as MedImg, which integrates diverse medical image datasets from multiple public sources and organizes all available data based on the organs and imaging modalities. Users can browse, retrieve, and download all images. Moreover, for each dataset, the platform provides its details and several samples to preview. MedImg database can be freely accessed at https://www.cuilab.cn/medimg/.

### Data collection and overview

This work aims at providing comprehensive medical images as possible for facilitating the researches on medical image analysis with deep learning. Considering the data privacy and ethics issue, we utilized the keywords of ‘medical image’ to obtain the medical image dataset with a public license. Ultimately, 103 out of 492 datasets meet the inclusion criterion and are downloaded, which are mainly derived from Grand Challenge and Kaggle, two websites holding AI-related competitions. Among them, 79 datasets include well-annotated labels, while not all medical image analysis requires the annotation. In addition, these datasets are built between 2007 and 2022, with the majority being established after 2015. This benefits the huge success of deep learning on image analysis, which, in turn, has led to the release of more image data.

These datasets cover various modalities harvested from including XR, computed tomography (CT), MRI, and other more. As shown in Figure 1A, the most frequently used modality in medical imaging is XR, constituting one-third of all data. XR is a more cost-effective and accessible form to visualize the internal structure of the body compared to CT and MRI. Other common data of medical imaging, such as histopathology slide (HS), endoscopy (ES), optical coherence tomography (OCT), electrocardiography (ECG), and ultrasound are also included. Moreover, the video and audio data of continuously monitoring diseases are kept as well. Figure 1B shows that lung, brain, skin, kidney, and liver are the mainly focused organs of medical images analysis. The diseases that frequently occur in these organs are varied precancerosis and cancers, which represents a leading cause of death [19] and, as such, motivates the growth of cancer-related images involving all body parts. On the other hand, numerous datasets containing images of the eye, heart, and knee are also available where imaging technologies are the primary means of lesion detection on these organs. For different deep learning tasks, the pipeline of preprocess, structure of model, and image annotation are totally distinct [20]. As depicted in Figure 1C, close to half of the datasets are employed for the image classification task, followed by segmentation accounting for 31%, which are the two most common tasks of deep learning applied on medical image analysis. Remaining data are separately designed to perform lesion detection, image registration, and regression. The number of images in different datasets ranges from 52 to 327,380, and the distribution of number of files in each dataset is exhibited in Figure 1D. A large dataset is essential for an excellent DL model [21]. Among these datasets, 35% contain over 5,000 images, 22 datasets of which hold more than 10,000 images. In sum, we incorporated a relatively comprehensive medical image repository with a coverage of 14 modalities, 9 organs, 1,622,956 images.

**Figure 1.**
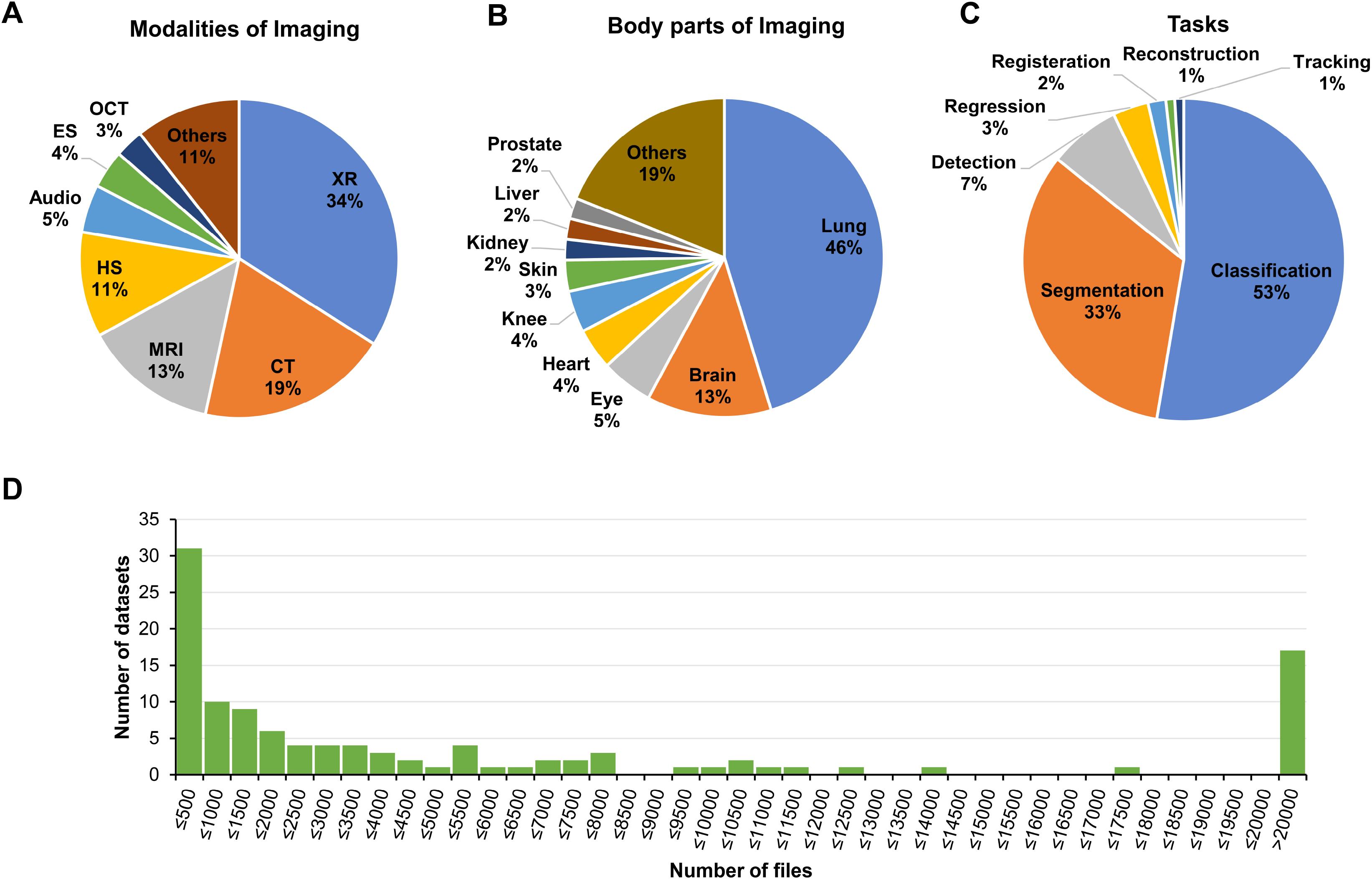
Summary of medical image datasets. (A) The percentage of imaging modalities of datasets. (B) The percentage of datasets across organs. (C) The percentage of deep learning tasks for all datasets. (D) The distribution of number of files in each dataset.

### Database implementation and utility

In order to provide researchers with an intuitive and efficient manner to access all the data, we establish an online database called MedImg, which stores and hierarchically organizes all medical images for users to quickly browse, query, and download. MedImg database (https://www.cuilab.cn/medimg/) is deployed based on Apache Tomcat server, of which the front end is implemented with HTML5 and CSS3; the interactive function and visualization are implemented with JQuery; the back-end is powered by Python Django framework. MedImg database can be accessed from multiple platforms such as PC and mobile phone without registration.

The MedImg online database is comprised of 4 main pages, including ‘Home’, ‘Search’, ‘Browse’, and ‘Download’ page. A brief introduction related to MedImg and its update information are contained in the ‘Home’ page. The navigation tree on the left side of the ‘Browse’ page is organized based on the data type (i.e. images, videos, and series), modalities of medical imaging (CT, MRI, and OCT etc.), and organs successively, as shown in Figure 2A. When the user clicks on a leaf of the navigation tree, the right side of this page presents the details and several representative samples of corresponding dataset (Figure 2B). The detail information includes the introduction for this dataset, the data format, sample size, organ source, and status of annotation. Figure 2B exhibits the preview result of different datasets with the format of images, audio, and video, respectively. Moreover, this dataset can be downloaded either from the detail panel or by clicking ‘Download Selected’ on the left navigation part. The ‘Search’ page allows users to quickly search for medical images of interest (Figure 2C). Users can directly input a keyword in the input box of ‘Dataset Name’. With an advanced search, users can retrieve the datasets through limiting multiple conditions, including the modalities of images, organ source of images, data format, with label or not, and task of the dataset. The retrieval result of real-time response is presented in the bottom of current page. Furthermore, all detailed information of datasets can be obtained by the ‘download’ icon button. In the search result, users can access the detail of this dataset by clicking on the dataset name link. It is noteworthy that users can separately download individual datasets or datasets of a specific class of their demand. Definitely, users have the option to download all data directly from the ‘Download’ page, although this might take some time due to the large size of data.

**Figure 2.**
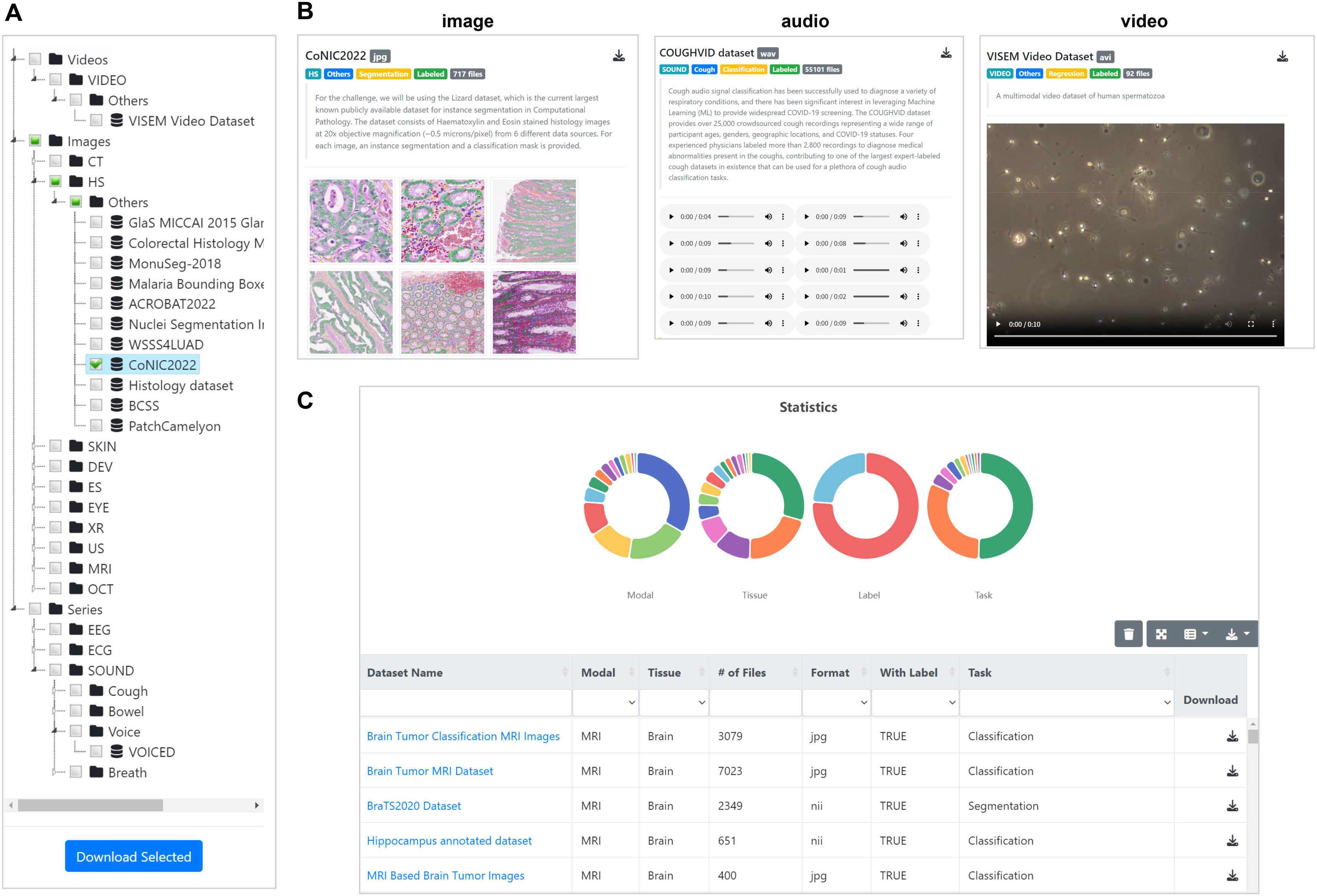
Overview of MedImg online database. (A) Navigation tree of the ‘Browse’ page. Users can download an individual dataset or a class of datasets via clicking on the ‘Download Selected’ button. (B) The details and preview for different types of datasets. (C) The main modules of search interface in MedImg. Users can input a keyword about the dataset or search combined with multiple conditions.

### Concluding remarks and outlook

The large and well-annotated datasets are the cornerstone of deep learning methods on medical image analysis. It is difficult to explore the generalization of an algorithm when using limited data, especially when it originates from a single community. At present, despite that the cost of manually labeling image data is high, more institutions and researchers have recognized the importance of well characterized datasets for improving deep learning algorithms and starts to publish and share their data. It is necessary to integrate those diverse resources and make them easily accessible. Therefore, we collected medical images across body parts and across modalities currently available and established a comprehensive online database, that is, MedImg. This database is an open-access and intuitive platform containing 103 datasets, more than 1.6 million images, which facilitates the related researchers to fast obtain benchmark datasets. MedImg is anticipated to contribute to driving the development of more generalized and robust deep learning-based algorithms for medical image analysis.

Undoubtedly, there still has several limitations. First, it is short of clinical information that are beneficial to the accuracy of AI-based diagnosis in current version. Second, it still needs to manually curate and normalize all datasets with a standard protocol for simplifying the preprocess step of users. We will continue to incorporate new medical image datasets to update the MedImg database.

## Authors’ contributions

**Bitao Zhong:** Data curation, Validation. **Rui Fan:** Software; Validation; Visualization. **Xiangwen Ji:** Investigation, Resources. **Qinghua Cui:** Conceptualization, Supervision, Writing – review & editing, Funding acquisition. **Chunmei Cui:** Conceptualization, Writing - original draft, Writing – review & editing, Funding acquisition.

## Acknowledgements

This study was supported by the grants from the National Natural Science Foundation of China (62025102, 81921001, 32301239), Scientific and Technological Research Project of Xinjiang Production and Construction Corps (2021AB028).

